# Cortical correlates of the auditory frequency-following and onset responses: EEG and fMRI evidence

**DOI:** 10.1101/076448

**Authors:** Emily B.J. Coffey, Gabriella Musacchia, Robert J. Zatorre

**Author notes:** **Corresponding author:** Emily Coffey, (alternative). **Conflicts of interest:** none.

## Abstract

The frequency following response (FFR) is a measure of the brain’s periodic sound encoding. It is of increasing importance for studying the human auditory nervous system due to numerous associations with auditory cognition and dysfunction. Although the FFR is widely interpreted as originating from brainstem nuclei, a recent study using magnetoencephalography (MEG) suggested that there is also a right-lateralized contribution from the auditory cortex at the fundamental frequency (Coffey et al., 2016c). Our objectives in the present work were to validate and better localize this result using a completely different neuroimaging modality, and document the relationships between the FFR and the onset response, and cortical activity. Using a combination of electroencephalography, fMRI, and diffusion-weighted imaging, we show that activity in the right auditory cortex is related to individual differences in FFR-f0 strength, a finding that was replicated with two independent stimulus sets, with and without acoustic energy at the fundamental frequency. We demonstrate a dissociation between this FFR-f0-sensitive response in the right and an area in left auditory cortex that is sensitive to individual differences in the timing of initial response to sound onset. Relationships to timing and their lateralization are supported by parallels in the microstructure of the underlying white matter, implicating a mechanism involving neural conduction efficiency. These data confirm that the FFR has a cortical contribution, and suggest ways in which auditory neuroscience may be advanced by connecting early sound representation to measures of higher-level sound processing and cognitive function.

**Significance Statement:** The frequency following response (FFR) is an electroencephalograph signal that is used to explore how the auditory system encodes temporal regularities in sound, and which is related to differences in auditory function between individuals. It is known that brainstem nuclei contribute to the FFR, but recent findings of an additional cortical source are more controversial. Here, we use functional MRI to validate and extend the prediction from magnetoencephalography data of a right auditory cortex contribution to the FFR. We also demonstrate a dissociation between FFR-related cortical activity from that related to the latency of the response to sound onset, which is found in left auditory cortex. The findings provide a clearer picture of cortical processes for analysis of sound features.

## Introduction

The Frequency Following Response (FFR) is an auditory signal recorded using electroencephalography (EEG) which offers a non-invasive view of behaviourally and clinically relevant individual differences in early sound processing (Krishnan, 2007; Skoe and Kraus, 2010; Kraus and White-Schwoch, 2015). Although the FFR itself is widely interpreted as having subcortical sources (Chandrasekaran and Kraus, 2010), its strength is correlatd with measures of cortical waves (Musacchia et al., 2008), and it is known to be modulated by cortical processes such as learning (Musacchia et al., 2007; Krishnan et al., 2008) and perhaps attention (Galbraith and Arroyo, 1993; Lehmann and Schönwiesner, 2014). Recent magnetoencephalography (MEG) evidence suggests that in addition to generators in brainstem nuclei, there is a direct contribution from the auditory cortex at the fundamental frequency (f0) with a rightward bias (Coffey et al., 2016c). However, MEG localization is indirect, relying on distributed source modeling to localize and separate cortical from subcortical sources, an approach whose limitations are still being explored (Attal and Schwartz, 2013). Validation of cortical involvement using more direct complementary methods is thus essential.

Features of the FFR vary between people, even within a neurologically normal young adult population (Hoormann et al., 1992; Ruggles et al., 2012; Coffey et al., 2016a). These differences have been linked to musical (Musacchia et al., 2007; Strait et al., 2009; Bidelman, 2013) and language (Wong et al., 2007) experience, and have been shown to be cognitively and behaviourally relevant, for example in the perception of speech in noise (Ruggles et al., 2012), consonance and dissonance (Bones et al., 2014), and in pitch perception bias (Coffey et al., 2016a). Similarly, the MEG FFR-f0 signal attributed to the right auditory cortex in our prior study was correlated with musical experience and fine frequency discrimination ability (Coffey et al., 2016c). These inter-individual variations provide a means of testing the hypothesis of an FFR-f0 contributor in the auditory cortex via fMRI: if stronger FFR-f0 encoding is partly indicative of greater phase-locked neuronal activity in the right auditory cortex, then FFR-f0 strength should be positively correlated with the magnitude of the BOLD response in the same area due to the increased metabolic requirements of this neural population (Magri et al., 2012). A related question concerns the generalizability of the MEG findings to other sounds. Our prior MEG finding relied on a synthetic speech syllable that produces a clear, consistent onset response and FFR (Johnson et al., 2005a; Skoe and Kraus, 2010). But the auditory system must also contend with sounds that include degraded or missing frequency information; to this end we used both the speech syllable and a piano tone without acoustic energy at f0.

As well as identifying FFR-f0–sensitive regions in the auditory cortex, it is useful to know if they can be dissociated from areas sensitive to other measures of early sound encoding, such as the timing of the transient onset response to sound, as suggested by behavioural dissociations (Johnson et al., 2005b; Kraus and Nicol, 2005; Skoe and Kraus, 2010). If this is the case, we would expect measures of the onset response and the FFR to correlate with BOLD activity in different cortical populations. To clarify timing-related results, we also obtained measures of white matter microstructure, which are related to signal transmission speed (Wozniak and Lim, 2006).

In the present study, we measured neural responses to two periodic sounds using EEG and fMRI, and assessed the relationships between measures of FFR-f0 strength, onset latency, and fMRI activity. Our primary aim was to test the hypothesis that individual differences in FFR-f0 strength is correlated with the magnitude of fMRI response in the right auditory cortex. We tested three additional hypotheses: that the FFR-f0 BOLD relationship is robust to stimuli with and without a fundamental; that an FFR-f0-sensitive area can be dissociated from an onset latency-sensitive area; and that timing-related results are correlated with the structure of the white matter directly underlying the auditory cortex.

## Materials and Methods

### Participants

We recruited 26 right-handed young adults divided into two groups: either musicians who practised at least one instrument regularly (>1.5hrs per week), or non-musicians with minimal exposure to musical training. All subjects reported having normal hearing and no neurological conditions and were compensated for their time. Normal or corrected-to-normal vision (Snellen Eye Chart) and pure-tone thresholds from 250 to 16kHz were measured to confirm sensory function (all but one subject had ≤ 20 dB HL pure-tone thresholds within the lower frequencies applicable to this study, 250–x2,000 Hz; this subject was included as stimuli are presented well above threshold binaurally and the opposite ear had a normal threshold). One subject was excluded due to a technical problem. The remaining 25 subjects (mean age: 25.8, SD: 5.0, 13 females) included 13 musicians and 12 non-musicians. Groups did not differ significantly in age (musicians mean: 25.2, SD = 5.6; non-musicians mean: 26.4, SD = 4.5; Wilcoxon rank sum test, two-tailed: Z = 0.84, p = 0.40) or sex (7 musicians and 6 non-musicians were female; Chi-square, two-tailed: X^2^(1,25) = 0.04, p = 0.85). Data about musical history were collected via an online survey (Montreal Music History Questionnaire; MMHQ (Coffey et al., 2011)). Musicians reported an average of 10,300 hours (SD: 5,000) of vocal and instrumental practice and training; 2 non-musicians reported ~400 hours of clarinet training as part of a school program, all others had no experience. The musicians varied in their instrument and musical style (main instruments: 3 keyboard, 2 woodwind, 9 strings including 5 guitar; main styles: 8 classical, 4 pop/rock, 1 traditional/folk). All experimental procedures were approved by the Montreal Neurological Institute Research Ethics Board.

### Study design

Subjects participated in separate EEG and MRI recording sessions on different days (randomized order; 13 subjects experienced the fMRI session first), during which they listened to blocks of repeated speech syllables or piano tones. Prior to the EEG session, subjects performed a set of computerized behavioural tasks (~30 mins), including fine frequency discrimination (reported below) and several other measures of musicianship and auditory system function (Nilsson, 1994; Foster and Zatorre, 2010) which relate to research questions that are not addressed here.

### Fine frequency discrimination assessment

Fine frequency discrimination thresholds were measured using a two-interval forced choice task and a 2-down 1-up rule to estimate the threshold at 79% correct point on the psychometric curve (Levitt, 1971). On each trial, two 250 ms pure sine tones were presented, separated by 600 ms of silence. In randomized order, one of the two tones was a 500 Hz reference pitch, and the other was higher by a percentage that started at 7 and was reduced by 1.25 after two correct responses or increased by 1.25 after an incorrect response. The task stopped after 15 reversals, and the geometric mean of the last 8 trials was recorded. The task was repeated 5 times, and the scores were averaged.

### Stimuli

We used two stimuli, a 100ms speech syllable (/da/) with a fundamental frequency of 98Hz that has been used extensively in previous studies as it elicits clear and replicable responses (Johnson et al., 2005a; Skoe and Kraus, 2010) (see Figure 1a,b, top), and a piano tone with the same nominal fundamental frequency and stimulus duration, but that had very little energy at the fundamental frequency (McGill University Master Samples database, Steinway piano G2 tone, right channel; (Opolko and Wapnick, 2006); see Figure 1c,d, top). In order to ensure that harmonic distortions created by the headphones did not reintroduce energy at the fundamental frequency (Norman-Haignere and McDermott, 2016), we measured sound output from both sets of earphones (S14, Sensimetrics Corp.; ER2, Etymotic Research) using a KEMAR Dummy-Head Microphone (GRAS, www.gras.dk), at the 80 dB SPL used in the experiment. Although the two earphones yielded slightly different amplitudes for each harmonic component, we found no evidence that energy had been reintroduced at the missing fundamental frequency.

**Figure 1.**
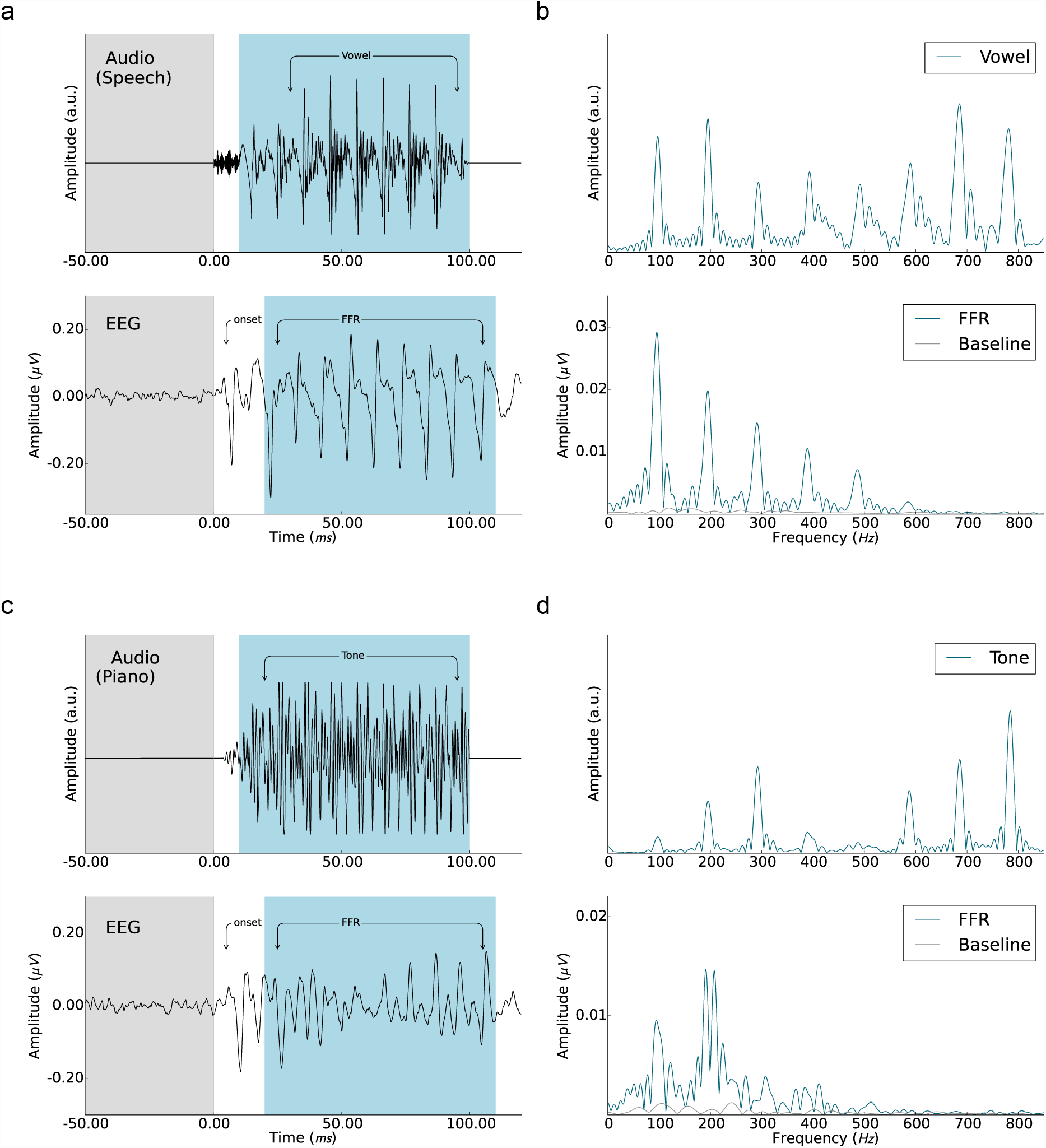
Auditory stimuli and averaged EEG responses. a) and b) show the speech stimulus (syllable: /da/, 98 Hz fundamental frequency) in the time and frequency domain (top) and the corresponding averaged responses isolated from the EEG recordings (bottom). c) and d) show the tone stimulus (piano ’G2’, 98 Hz missing fundamental frequency) in the time and frequency domain (top) and the EEG responses (bottom). The prestimulus baseline (-50 to 0ms) and the frequency-following response (FFR) periods (20 to 110ms after sound onset) are marked in grey and blue, respectively.

### fMRI data acquisition

The stimulation paradigm took into account the constraints of each of the imaging modalities such that almost identical versions could be presented during the independent EEG and BOLD fMRI recording sessions. Each interval between scans, defined as a block, comprised a series of 20 stimuli of the same type (inter-stimulus interval: ~200 ms, jittered by 0-10 ms, randomized) as well as silent breaks (Figure 2), which were included to reduce the effects of repetition suppression and enhancement that can differ between people (Chandrasekaran et al., 2012). Stimuli were presented binaurally at 80 dB ± 1dB SPL, using a custom-written script (Presentation, Neurobehavioral Systems, Albany, CA, USA), using MRI-compatible headphones (S14, Sensimetrics Corp.) via foam inserts placed inside the ear canal. Auditory stimulation was timed so as to maximize the hemodynamic response during fMRI recording to sound during the subsequent acquisition (i.e. ∼5-7 sec after the onset of the stimulus block), but its exact timing was jittered (0-1s, randomized) so as to reduce confounds with periodic sources of noise and of top-down expectations. Speech or Piano tone blocks were presented pseudorandomly, along with Relative Silence baseline blocks (for a total of 120 syllable volumes, 120 tone volumes, and 90 baseline volumes). Subjects were asked to listen actively for oddball stimuli (80% normal amplitude) and indicate via button press (right index and middle finger) during the scan following stimulation if one had occurred or not. Oddballs were present in 30% of the blocks and replaced one of the last 4 stimuli in a block. To control for preparatory motor activity associated with button pressing, baseline volumes included a single stimulus ∼1-2 seconds from the end of the block to which subjects responded during the scan with a button press. Nine subjects experienced a slight experimental variation in which the single stimulus was presented ∼4 seconds from the end of the block; this difference was controlled for in each GLM model.

**Figure 2.**
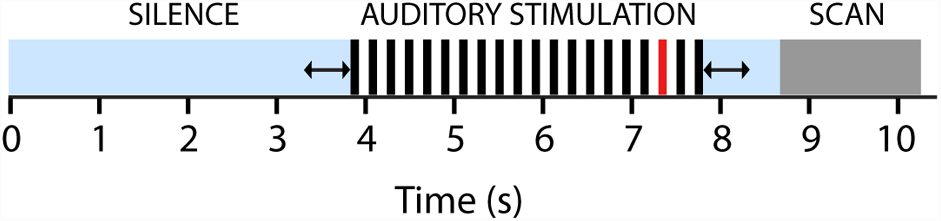
Auditory stimulation paradigm. Each stimulation block consisted of twenty repetitions of the same stimulus (either speech or piano), which was situated within a period of silence and jittered to minimize physiological confounds (see Methods for details). The same design was used for the EEG and fMRI recording sessions. In 30% of blocks, a quieter stimulus was presented in place of one of the last 4 stimuli (indicated in red). Subjects were asked to indicate whether there had been an oddball after each block, in order to control for attention.

fMRI data were acquired using EPI whole head coverage on a Siemens 3 Tesla scanner with a 32-channel head coil (Siemens Trio, Erlangen, Germany) at the McConnell Brain Imaging Center at the Montreal Neurological Institute using a sparse sampling fMRI paradigm (Belin et al., 1999; Hall et al., 1999); which avoids confounding the BOLD signal of interest with effects due to loud noise from gradient switching (voxel size 3.4mm^3^, 42 slices, TE 49 ms, TR ~10210ms). We implemented a cardiac gating procedure such that each scan was triggered by the cardiac cycle following the stimulation block (Guimaraes et al., 1998) in order to address research questions that are not reported here. This resulted in an average block length difference as compared with EEG of ~500ms, and total fMRI scan time was approximately 1hr (3 runs of 19 mins each). To reduce subject fatigue, anatomical MRI scans were acquired between fMRI runs, during which subjects were instructed to lie still and rest.

### FMRI analysis

FMRI data were analysed using FSL software (fMRIB, Oxford, UK) (Smith et al., 2004; Jenkinson et al., 2012). Images were motion-corrected, b0 unwarped and registered to the T1-weighted anatomical image using boundary-based registration (Greve and Fischl, 2009), and spatially smoothed (5mm FWHM). Each subjects’ anatomical image was registered to MNI 2mm standard space (12-parameter linear transformation). For 6 subjects, gradient field maps had not been acquired; these were substituted by an average of the other 19 subjects’ gradient field maps in standard space, transformed to native space (12-parameter linear transformation). Task-related BOLD responses of each run were analyzed within GLM (FEAT; (Beckmann et al., 2003)), including 3 conditions (Relative Silence, Speech, Piano). For each scan, contrast images were computed for Speech > Relative Silence and Piano > Relative Silence, and three runs per subject were combined in a fixed-effects model. Within and between-group analyses were performed using random effects models in MNI space (FLAME 1 in FSL; the automatic outlier deweighting option was selected). In order to test the specific hypotheses of interest and to localize areas of sensitivity to FFR-f0 strength within the auditory cortex, a bilateral auditory cortex region of interest (ROI) was defined using the Harvard-Oxford cortical and subcortical structural atlases implemented in FSL: regions with a probability greater or equal to 0.3 of being identified as Heschl’s gyrus (HG) or planum temporale (PT) were included, and the resulting ROI was dilated by two voxels to ensure that the central peaks of the cortical signal generators found in previous work (Coffey et al., 2016c) were well within the ROI.

To evaluate the main research questions, we entered the FFR-f0 and wave A latency values into a whole-sample GLM model, separately for the Speech vs. Relative Silence and Piano vs. Relative Silence contrast (the minor difference in the Silent blocks described above was entered as a covariate of no interest). For multiple comparisons correction for each research question, we applied voxel-wise correction as implemented in FEAT (Gaussian Random Field-theory-based, p <0.05 one-tailed; statistical maps thresholded above Z = 2.3 are presented in Figures 3 and 4; significant clusters are reported in the text). To gain additional evidence that the brain area identified as being sensitive to FFR-f0 strength in the Speech condition was also related to FFR-f0 strength in the Piano condition, we ran a conjunction analysis in which the Piano condition regression analysis was masked by the significant result from the speech regression analysis. For further analysis of relationships to fine frequency discrimination threshold and musicianship, we extracted a measure of BOLD activity (mean percent change of parameter estimate) from the small cortical areas that were found to be significantly related to FFR-f0 strength, in each contrast.

**Figure 3.**
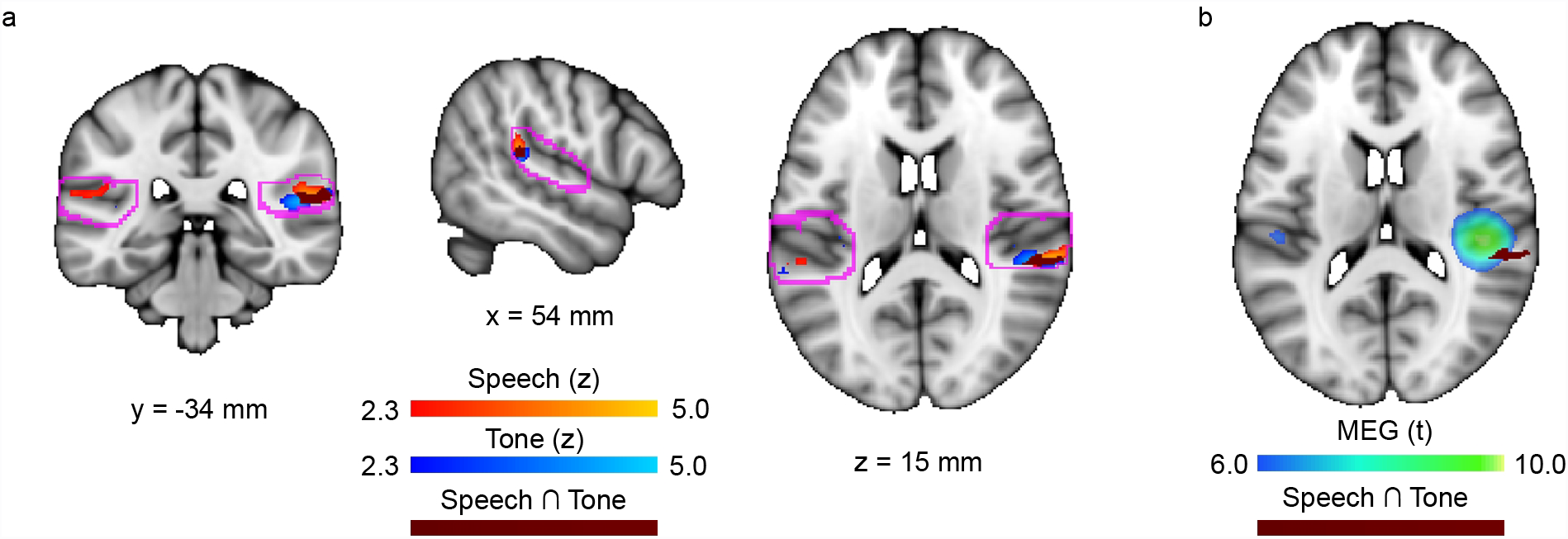
Areas within the auditory cortex that are sensitive to FFR-f0 strength. a) Coronal, sagittal, and horizontal brain slices showing statistical maps where BOLD signal was related to FFR-f0 strength for each stimulus set; greater FFR strength was related to higher BOLD signal in the right planum temporale (Speech: orange; Piano: blue). Overlapping regions are indicated in maroon. Bilateral regions of interest encompassing the auditory cortex bilaterally are delineated in pink. b) A horizontal slice showing the location of FFR-f0 sensitive cortex in both conditions (maroon) in relation to the previous result of a right auditory cortex contribution to the FFR-f0, from MEG (Coffey et al., 2016c); note that fMRI and MEG differ in their spatial resolution.

**Figure 4.**
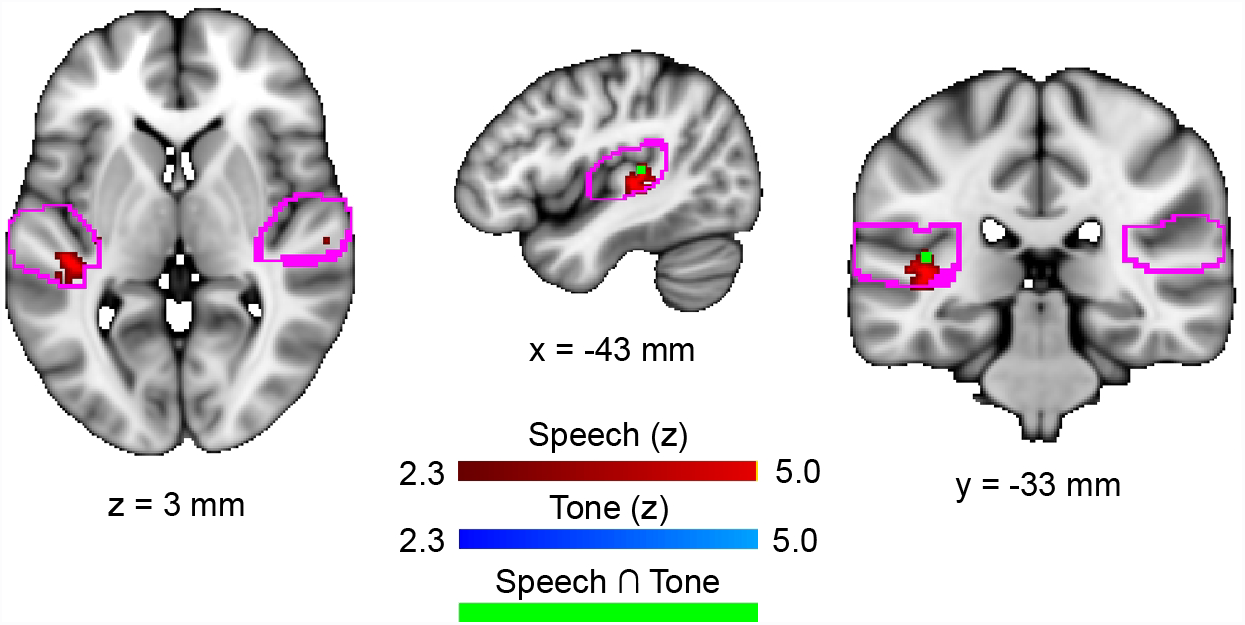
Areas within the auditory cortex that are sensitive to onset latency. Horizontal, sagittal, and coronal slices showing statistical maps where BOLD activity was correlated with the latency of the onset response in the Speech condition (red). Shorter latencies were related to lower BOLD signal in left Heschl’s sulcus. No significant areas were found in the Piano condition, though a sub-threshold cluster was observed to overlap with the Speech result (Z = 2.35; visible in green).

### EEG acquisition

Because the EEG version of the paradigm did not include baseline silent blocks nor cardiac gating, the total recording time was 45 minutes. This was split into three parts, between which short breaks were given. During the recording, subjects sat comfortably in a magnetically shielded room. A Biosemi active electrode system (ActiveTwo) sampled at 16kHz was used to to record EEG from position Cz, with two earlobe references, and grounds placed on the forehead above the right eyebrow. Stimuli were presented using a custom-written script (Presentation (Neurobehavioral Systems, Albany, CA, USA), delivered binaurally via insert earphones (ER2, Etymotic Research, www.etymotic.com). Each stimulus was presented 2400 times, at alternating polarities to enable cancelling of the cochlear microphonic (Skoe and Kraus, 2010). We recorded stimulus onset markers from the stimulus computer along with the EEG data via parallel port. Subjects were asked to keep their bodies and eyes relaxed and still during recordings, and were provided with a small picture affixed to the wall as a reminder.

### EEG analysis

Data analysis was performed using the EEGLAB toolbox (v13.5.4b (Delorme and Makeig, 2004)), the ERPLAB plugin (v5.0.0.0), and custom MatLab scripts (MATLAB 7.12.0, The MathWorks Inc., Natick, MA, 2000). Each recording was band-pass filtered (80-2000Hz; Butterworth 4^th^ order, zero-phase, as implemented in EEGLAB), epoched (-50 to 200ms around the onset marker), and DC correction was applied to the baseline period. Fifteen percent of epochs having the greatest amplitude were discarded for each subject; this served to remove the majority of epochs contaminated by myogenic activity (confirmed by inspection), yet retain equal numbers of epochs per subject for the computation of phase-locking value (PLV): a measure of FFR strength that is highly correlated with spectral amplitude but that is more statistically sensitive (Zhu et al., 2013). For each subject and stimulus type, a set of 400 epochs from the total pool (2040) was selected randomly with replacement. Each epoch was trimmed to the FFR period (20-110ms after sound onset), windowed (5 ms raised cosine ramp), zero-padded to 1 s to allow for a 1 Hz frequency resolution, and the phase of each epoch was calculated by discrete Fourier transform. The PLV for each epoch was computed by normalizing the complex discrete Fourier transform by its own magnitude and averaging across 1000 iterations. Mean f0 strength was taken to be the mean PLV at f0 (peak +/− 2 Hz), for each subject and stimulus (see ’Appendix: analysis methods’ item 5, in (Zhu et al., 2013) for formulae).

To obtain onset latency, epochs were averaged together by polarity to correct for any effect of the cochlear microphonic (i.e. negative, positive; (Wever and Bray, 1930)) and summed to form the time domain average. To select an onset peak for analysis, we generated a grand average for each stimulus across all subjects, and compared individual waveforms with it, as suggested in (Skoe and Kraus, 2010); we selected wave A for further analysis for replicability across subjects. An experienced rater who was blind to subject identity and group selected wave A peak latencies for each subject and condition by visual inspection. These were confirmed by a custom automatic algorithm (Spearman’s correlation between the manually and automatically selected wave A latency for the speech stimulus: r_s_ = 0.98, p <0.001; piano stimulus: r_s_ = 0.85, p <0.001). Manually selected latencies were deemed to be similar yet were preferred, as it was sometimes necessary for the less clear piano onset to select between two local peaks.

Distributions of FFR-derived measures frequently fail tests of normality, as is the case here: we performed Kolmogorov-Smirnov tests on the FFR-f0 and wave A latency for each condition, and in each case rejected the hypothesis of a normal distribution (p <0.05). Non-parametric statistics were therefore used unless otherwise specified. We compared FFR-f0 and wave A latency across musicians and nonmusicians using one-tailed Wilcoxon rank sum tests, and assessed correlations between start age and total practice hours, and FFR-f0 strength and wave A latency using Spearman’s rho; r_s_.

### Anatomical data

Between the first and second functional imaging run, we recorded whole-head anatomical T1-weighted images (MPRAGE, voxel size 1mm3). Freesurfer was used to automatically segment each brain (Fischl et al., 2002). Between the second and third run, we recorded diffusion-weighted images (DWI; 99 directions, voxel size 2.0mm^3^, 72 slices, TE 88 ms, TR 9340 ms, b = 1000s/mm2). Diffusion-weighted images were corrected for eddy current distortions, brains were extracted from unweighted images, and a diffusion tensor model was fit using FSL’s ’dtifit’ function to obtain voxelwise maps of the diffusion parameters (fractional anisotropy, mean diffusivity, and axial diffusivity). Radial diffusivity was calculated as the mean of the second and third eigenvalues of the diffusion tensor.

Regions of interest below the grey matter that were identified as Heschl’s gyrus and sulcus by Freesurfer segementation (Destrieux et al., 2010)) were created for each hemisphere by transforming surface labels from each participant’s native space into their diffusion-weighted volume space, projecting them to a depth of 2mm (parallel to the cortical surface), and visually confirming that voxels lay in white matter for each participant; these masks are used to address questions of lateralization and relationships to fMRI and EEG results in white matter that is most directly related to the auditory cortex (Shiell and Zatorre, 2016). Transformation matrices were calculated between DWI space and structural space (T1-weighted image, FLIRT, 6 degrees of freedom) and to a 1mm FA template (FMRIB58_FA_1mm, FLIRT, 12 degrees of freedom), concatenated, and their inverses used to transform individual Heschl’s gyrus and sulcus masks to diffusion space to extract diffusion measures.

To address research questions about possible differences in the microstructure of white matter underlying regions of the auditory cortex that were found to be sensitive to onset timing, we first evaluated correlations between onset latency in the Speech condition and two measures of white matter microstructure, fractional anisotropy (FA) and mean diffusivity (MD), in each white matter region of interest (corrected for multiple comparisons, alpha = 0.05/4). To better understand the mean diffusivity result, we also assessed correlations between onset latency in the Speech condition and sub-components of mean diffusivity: axial diffusivity (AD) and radial diffusivity (RD). To assess the lateralization of the observed mean diffusivity finding, we statistically compared the correlations in each auditory cortex using Fisher’s r-to-Z transformation (Steiger, 1980). Finally, we predicted a negative correlation between BOLD response and MD values in the left auditory cortex based on the BOLD- onset and onset-MD correlations, and tested this relationship for statistical significance using Spearman’s rho (one-tailed).

## Results

### Attention control

Subjects correctly identified most of the blocks as either containing oddball (quieter) stimuli or not during both sessions (EEG mean accuracy = 85.3%, SD = 11.2; fMRI mean accuracy = 90.9%, SD = 8.7); this served to confirm that subjects were attending to the stimuli.

## Regression of FFR-f0 with BOLD fMRI data

### Speech condition

In the Speech > Relative Silence contrast, FFR-f0 strength was significantly correlated with BOLD signal in the right (but not left) posterior auditory cortex / planum temporale (Fig. 3a,b; the significant cluster has a volume of 128 mm3 and is centred at: x = 60, y = −34, z = 14 mm; 2mm MNI152 space; Z = 3.99). Musicians showed significantly stronger BOLD responses than non-musicians within the region identified as being significantly sensitive to FFR-f0 strength (Wilcoxon rank sum test, one-tailed: Z = 2.15, p = 0.016; musician mean: 0.53% change of parameter estimate (SD = 0.50); non-musician mean: 0.16% (SD = 0.47)), although the between-group differences in FFR-f0 strength did not reach significance (Z = 0.24, p = 0.4; musician mean PLV:

0.14 (SD = 0.06); non-musician mean PLV: 0.12 (SD = 0.04).

### Piano condition

In the Piano > Relative Silence contrast, FFR-f0 was significantly correlated with BOLD signal in the right AC region (the significant cluster has a volume of 112 mm3 and is centred at: x = 52, y = −34, z =12 mm; 2mm MNI152 space; Z = 4.10; Fig. 3). The conjunction analysis revealed that the majority of the region identified as sensitive to FFR-f0 in the Speech condition was also significantly related to FFR-f0 in the Piano condition (i.e. 112 mm3 out of 128 mm3). As in the Speech condition, musicians showed significantly stronger BOLD responses than non-musicians within the region identified as being significantly sensitive to FFR-f0 strength (Wilcoxon rank sum test: Z = 2.15, p = 0.016; musician mean: 0.37% (SD =0.54); non-musician mean: 0.07% (SD = 0.44)). The between-group differences in FFR-f0 strength did not reach significance, although a trend was suggested (Z = 1.50, p= 0.067; musician mean PLV: 0.09 (SD = 0.04); non-musician mean PLV: 0.07 (SD = 0.02).

In addition to the right AC area, several voxels within the left hemisphere ROI at the extreme anterior end were found to be significantly related to FFR-f0 strength. This region does not overlap with the left auditory cortex FFR-f0 generator derived from the MEG, nor does it appear to be in homologous regions the the right auditory cortex finding, but for completeness we explored this finding by inspecting the statistical maps from each condition in the vicinity of the ROI borders. The left anterior cluster was located in the posterior division of the superior temporal sulcus (centre: x= −66, y −18, z= −2 mm; 2mm MNI152 standard brain; Z = 4.1). A similar cluster was also present in the Speech vs. Relative Silence condition (maximum: x= −58, y −16, z= −4 mm, Z = 2.73). One additional cluster was found in the left posterior parietal operculum, outside of the ROI (Piano > Relative Silence condition: x=-46, y=-40, z=24 mm; Z = 2.78; Speech > Relative Silence condition: x=-48, y=- 40, z=26 mm; Z = 3.52). Neither of these clusters appeared in the right hemisphere homologue structures, nor did there appear to be other f0-sensitive clusters near the right hemisphere ROI borders.

## Regression of onset latency with BOLD fMRI data

### Speech condition

In the Speech > Relative Silence contrast, longer wave A latencies were correlated with greater BOLD signal in the left (but not right) Heschl’s sulcus (Fig. 4; the significant cluster has a volume of 40 mm3 and is centred at: x = −42, y = −32, z =6 mm; 2mm MNI152 space; Z = 4.29; Fig. 5 a,c). BOLD signal was not significantly related to shorter latencies, which are considered to index better functioning, in any regions. We did not observe a difference between musicians and non-musicians in BOLD response within the area sensitive to wave A latency (Wilcoxon rank sum test: Z = −0.73, p = 0.46), nor in the wave A latency values (Z = −0.41, p = 0.34).

**Figure 5.**
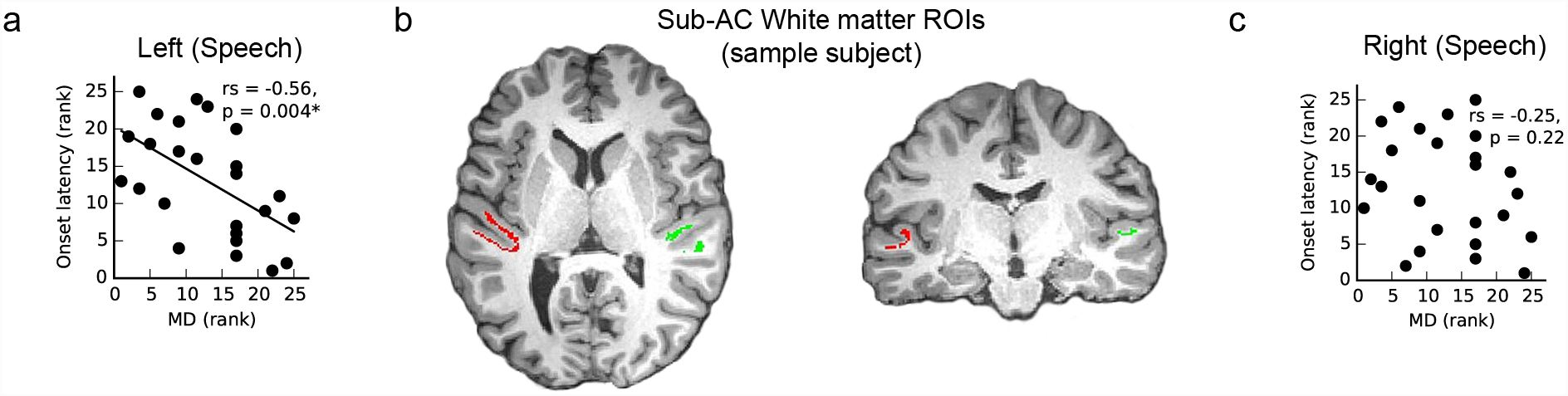
White matter microstructure is related to onset latency. a) Left hemisphere mean diffusivity values within anatomically defined regions of interest show a significant correlation: shorter latencies were related to greater mean diffusivity values. b) White matter regions of interest underlying auditory cortex on a single example subject overlaid on a T1-weighted anatomical image, for illustrative purposes. c) Similar analyses on the right side did not show any significant correlation. This analysis was carried out only in the Speech condition, as onset latencies to the natural piano tone were more variable (see Methods and Results for details).

### Piano condition

No areas were significantly related to piano wave A onset latency in the Piano > Relative Silence contrast. Although a sub-threshold peak was observed within the area sensitive to latency in the Speech condition (x = −42, y = −32, z = 10; Z=2.35; 2mm MNI152; see the conjunction (green) in Fig. 4 for location), we carry out secondary analyses relating to onset latency only in the significant Speech condition.

## Onset latency and microstructure of white matter underlying auditory cortex

Onset latency in the Speech condition was significantly correlated with average MD values within the white matter ROI underlying Heschl’s gyrus and sulcus in the left hemisphere (two-tailed, corrected for multiple comparisons, alpha = 0.05/4; r_s_ = −0.56, p = 0.004; Fig. 5a), but not in the right hemisphere (r_s_ = −0.25, p = 0.22; Fig. 5c). The correlation between onset latency and MD was significantly greater in the left than the right hemisphere (Fisher’s r-to-z transformation, one-tailed, Z = 1.765, p = 0.039). Significant relationships between onset latency and mean FA were not observed in the left hemisphere ROI (r_s_ = −0.19, p = 0.36), nor right hemisphere ROI (r_s_ = −0.10, p = 0.62).

Both axial diffusivity and radial diffusivity showed similar patterns in their relationships to onset latency as did mean diffusivity in the left hemisphere (AD vs. onset: r_s_ = −0.62, p = 0.0008; RD vs. onset latency: r_s_ = −0.45, p = 0.024), and no significant relationship in both cases in the right hemisphere (AD vs. onset: r_s_ = −0.24, p = 0.25; RD vs. onset latency: r_s_ = −0.22, p = 0.29).

If a greater BOLD response and lower MD are both indices of neural conduction inefficiency, then we would predict a negative correlation between MD under left Heschl’s gyrus and BOLD response in the overlying grey matter. This is in fact the case (r_s_ = −0.42, p = 0.019). Musicians did not differ significantly from non-musicians in MD on either side (reported values are two-tailed; left: Z = 0, p = 1.0; right: Z = 0.14, p = 0.89).

### Fine frequency discrimination

#### Assessment of fine frequency discrimination skills

The mean fine frequency discrimination threshold (FF) was 1.40% overall, (SD = 1.45). Musicians had lower fine frequency discrimination thresholds than non-musicians as expected (musician mean: 0.58%, SD = 0.37; non-musician mean: 2.29%, SD = 1.66; Z = 3.40, p < 0.001). The BOLD signal strength extracted from the FFR-f0 sensitive region was not significantly correlated with fine frequency discrimination (reported p-values are one-tailed; Speech condition: r_s_ = −0.18, p = 0.19; Piano condition: r_s_ = −0.13, p = 0.27) and nor was frequency discrimination and BOLD signal significantly related within the FFR-f0-sensitive auditory cortex regions (Speech condition: r_s_ = −0.31, p = 0.06; Piano condition: r_s_ = −0.24, p = 0.12).

### Discussion

Our results demonstrate that hemodynamic activity in the right posterior auditory cortex is sensitive to FFR-f0 strength, a finding that was replicated in two separate stimulus sets with and without energy at the fundamental frequency, and which conforms to predictions arising from our prior MEG study (Coffey et al., 2016c). The right-lateralized FFR-f0-sensitive region was dissociable from a left-lateralized region in Heschl’s sulcus that was sensitive to the latency of the onset response. This finding was further supported by a significant relationship between onset latency and the microstructure of the white matter immediately underlying primary auditory areas in the left (but not right) hemisphere, and a significant correlation between BOLD response in the onset-sensitive region and mean diffusivity in underlying white matter. A lateralization of the relationship between onset timing and white matter microstructure is supported by a direct comparison of correlation strength.

#### Relationship between BOLD-fMRI and FFR-f0

Our primary aim was to adduce evidence in favour of a cortical source for the FFR (Musacchia et al., 2008; Coffey et al., 2016c), to which end we tested the hypothesis that the FFR-f0 strength is correlated with fMRI signal in the right auditory cortex. We reasoned that if inter-individual variations in FFR-f0 strength reflect differences in the coherence or number of phase-locked neurons within this population, these variations should be paralleled by differences in localized metabolic requirements that would manifest as an FFR-f0 sensitive area in the fMRI signal. This hypothesis was supported, and further corroborates preliminary reports of an FFR-like signal measured intracranially from the auditory cortex (Bellier et al., 2014). Together with previous MEG work (Coffey et al., 2016c), our data suggest that findings based on the FFR-f0 should not be assumed to have purely brainstem origins. Because these findings are in agreement with the conclusion based on MEG data that there is a cortical component to the FFR, it also supports the use of the new MEG-FFR method to observe the sources of the more commonly used scalp-recorded EEG-FFR.

That two independent stimuli result in overlapping areas of FFR-f0 sensitivity, whether f0 energy is present in the auditory signal or not, suggests that the sound representation within this region may be involved in computation of pitch at an abstract level. Missing fundamental stimuli are known to produce FFRs with energy at the fundamental frequency (Smith et al., 1978; Galbraith, 1994), and inter-individual variability in f0 strength is related to inter-individual variability and conscious control of missing fundamental perception, though not in a linear manner (Coffey et al., 2016b). Together, these results raise the possibility that top-down task modulation and perhaps experience-related modulation of FFR-f0 strength observed previously (e.g. (Musacchia et al., 2007; Lehmann and Schönwiesner, 2014)) could be occurring at the level of the auditory cortex, although it does not rule out the possibility that the strength of sub-cortical FFR-f0 components are also modulated concurrently. The right auditory cortex has been implicated previously in missing fundamental pitch computation (Schneider and Wengenroth, 2009): patients with right temporal-lobe excisions that include the right lateral auditory cortex have difficulty perceiving the missing fundamental (Zatorre, 1988), and asymmetry in grey matter volume in lateral Heschl’s gyrus is related to pitch perception bias (Patel and Balaban, 2001; Schneider et al., 2005). While the FFR-f0 is likely not a direct representation of pitch (Gockel et al., 2011), our results further connect the FFR’s pitch-bearing information to processes taking place in auditory cortex regions that represent pitch in an invariant fashion (Penagos, 2004; Bendor and Wang, 2006; Norman-Haignere et al., 2013).

#### Relationship between BOLD-fMRI and onset response latency

The onset response and the FFR-f0 may be represented in different auditory streams (Kraus and Nicol, 2005), as each measure co-varies with distinct behavioural and clinical measures (Kraus and Nicol, 2005; Skoe and Kraus, 2010); we therefore wanted to test for a dissociation in the cortical areas sensitive to each measure. However, the mechanistic basis for predicting a greater fMRI signal with a greater amplitude (as in the FFR-f0 analysis) does not hold true for latencies; we do not expect shorter onset latencies to necessarily relate to a larger population of neurons firing and therefore greater metabolic requirements that would be reflected in the BOLD signal, nor could onset-related sensitivity be directly related to the generation of the onset response, which occurs in the brainstem before sufficient time has elapsed for neural transmission to the cortex (Parkkonen et al., 2009). We therefore tested both positive and negative relationships. We found only a significant negative relationship: greater BOLD responses are related to longer latencies in left auditory cortex.

In order to confirm this result and partly inform a mechanistic explanation, we investigated the microstructure of white matter in regions of interest directly underlying Heschl’s gyrus and sulcus. In a study of the relations between task-related BOLD signal in human grey matter and measures of white matter microstructure, Burzynska et al. reported that greater microstructural integrity of major white matter tracts was negatively related to BOLD signal, which was interpreted as better quality of structural connections allowing for more efficient use of cortical resources (Burzynska et al., 2013). If a similar mechanism is at work here, we would expect that the BOLD sensitivity to onset latency should be paralleled by a relationship between WM microstructure and onset latency, and this relationship should also show a left lateralization. We confirmed these relationships in the mean diffusivity measure (corroborated in radial and axial diffusivity sub-components) but not the fractional anisotropy measure. FA is a measure of relative degree of sphericity vs. linearity of the diffusion tensor, which may not be as relevant a measure in white matter underlying GM as in major white matter tracts, due to the presence of association fibres. Although the nature of the observed structural sensitivity to onset latency in the white matter at the cellular level cannot be ascertained from diffusion-weighted data, the direction of the observed relationships between onset latency, BOLD signal, and diffusivity suggests that lower mean diffusivity in white matter and lower BOLD response in overlying areas are associated with greater neural conduction efficiency within the ascending white matter pathways that carry the onset signal to the cortex. Further work is needed to confirm the white matter finding reported here and to clarify whether it reflects more extensive white matter differences throughout the ascending auditory pathway, as would be predicted by the relationship to the timing of the subcortically-generated onset response.

#### Relative lateralization

We found a right-lateralized relationship between BOLD signal and FFR-f0, and a left-lateralized relationship between BOLD signal and onset latency (which was supported by a lateralization in underlying white-matter structure). Our results are in agreement with previous evidence of a relative specialization of the right AC for aspects of pitch and tonal processing (Zatorre, 1988; Zatorre and Belin, 2001; Patterson et al., 2002; Hyde et al., 2008; Mathys et al., 2010; Albouy et al., 2013; Herholz et al., 2015; Matsushita et al., 2015; Cha et al., 2016).

There is also experimental evidence for a complementary left AC specialization for aspects of temporal resolution (reviewed in (Zatorre et al., 2002; Poeppel, 2003; Wong et al., 2008)), although the interpretation of such findings and how they relate to linguistic processes is controversial (Scott and McGettigan, 2013). Nonetheless, the pattern of results reported here, particularly that onset response timing is related to both BOLD response in primary auditory cortex grey matter and in the structural properties of underlying white matter in the left but not right hemisphere, does favour the proposal of a relative specialization for enhanced temporal resolution in the left auditory cortex. Further work is needed to determine where in lower levels of the auditory system this lateralization first emerges.

#### Relationship to training and behaviour

We found that BOLD signal was significantly greater in musicians for both stimuli, in accord with several prior studies (Pantev and Herholz, 2011), and likely reflecting enhanced processing of pitch information. We found significant effects of musician training in the fMRI data. Although the FFR-f0 effects do not reach significance, differences have not been consistently observed in similar sample sizes (Musacchia et al., 2007; Wong et al., 2007; Lee et al., 2009; Strait et al., 2012), possibly because they may be eclipsed by large inter-individual variations (Coffey et al., 2016b). Previous work also showed clearer behavioural relationships to FFR-f0 components that had been separated by their source using MEG than to the FFR-f0 strength measured with EEG (Coffey et al., 2016c); it is therefore possible that the compound nature of the EEG signal obscures behavioual relationships of interest here.

### Conclusion

Our results validate and extend the prediction from magnetoencephalography data of a right auditory cortex contribution to the FFR and show a dissociation in early cortical auditory regions of the FFR-f0 and onset timing, providing further evidence that the auditory cortex is both functionally and structurally lateralized. The finding that inter-individual differences in FFR strength and onset latency in a population of normal-hearing young adults have cortical correlates supports the idea that these measures represent variations in input quality to different higher-level cortical functions and processing streams, which in turn influences perception and behaviour.

